# Exploring new nanopore sensors from the aerolysin family

**DOI:** 10.1101/2024.04.07.588449

**Authors:** Nuria Cirauqui, Juan F. Bada Juarez, Fernando Augusto T. P. Meireles, Julian Barry, Monika Bokori-Brown, Maria J. Marcaida, Chan Cao, Matteo Dal Peraro

**Affiliations:** Institute of Bioengineering, School of Life Sciences, EPFL, Lausanne 1015, Switzerland; University of Exeter, Exeter EX4 4QD, United Kingdom; Department of Inorganic and Analytical Chemistry, Chemistry and Biochemistry, University of Geneva, Quai Ernest-Ansermet 30, CH-1211 Geneva, Switzerland

## Abstract

Aerolysin-like proteins are a family of β-pore-forming toxins which are widely present in all kingdoms of life. Recently, this family of proteins is gaining attention because of their biotechnological application as nanopore sensors for sensing and sequencing of biomolecules. Here, we explore the possibilities of using the knowledge of the sequence and structure of proteins to screen and explore new potential nanopore candidates. However, in spite of the conserved structural fold, the sequence identity in this family is very low. This complicates their sequence alignment, hindering the understanding of their pore structure and properties, therefore limiting further biotechnological applications. In an attempt to further understand the properties of aerolysin-like pores, we analyzed the pore structure of three family members, *Clostridium perfringens* epsilon toxin (ETX), *Laetiporus sulphureus* lectin (LSL) and *Bacillus thuringiensis* parasporin-2, comparing it to aerolysin. Their structure and sensing capabilities for ssDNA were first assessed by *in silico* methods. Moreover, ETX was characterized experimentally in planar lipid membranes for the detection of biomolecules. We found that ETX can form three distinct pore conformations, each presenting a specific open pore current, and only one of them being able to translocate ssDNA. When the ssDNA translocate through ETX, the depth of current blockage is higher compared to aerolysin which indicates a higher sensitivity for molecular sensing. Our findings open a new venue for improving and diversifying nanopore capabilities for molecular sensing.

## Introduction

Aerolysin-like proteins are a sub-family of β-pore forming toxins that are present in all forms of life, from prokaryotic to eukaryotic organisms^1,2^. They all share in their monomeric state the same fold as aerolysin, which is the first and best structurally characterized protein among these toxins (**Figure 1A**). Moreover, based on their structural similarity they are thought to present similar pore-forming mechanisms and pore architecture^2^. These toxins are secreted as soluble monomers and after oligomerization into a pre-pore state bind to the membrane, inserting the stem loop (cyan in **Figure 1A** and **Supplementary Figure S1**) into the lipid bilayer, followed by strands β2-3 (orange and purple in **Figure 1A** and **Supplementary Figure S1**), to finally form a β-barrel pore. Upon insertion, there is a sliding of contacts on the stem loop amino acids, forming both the transmembrane barrel and the rivet loop^3^. The pore-formation can be driven by an arrangement of the double β-barrel fold present already in the pre-pore oligomer but that reaches its full stability only at the mature pore conformation (closeup in **Figure 1A** and **Supplementary Figure S1**)^2^. This novel fold characterizing the pore family is thought to be responsible for the ultra-stability of aerolysin pores^2^.

**Figure 1.**
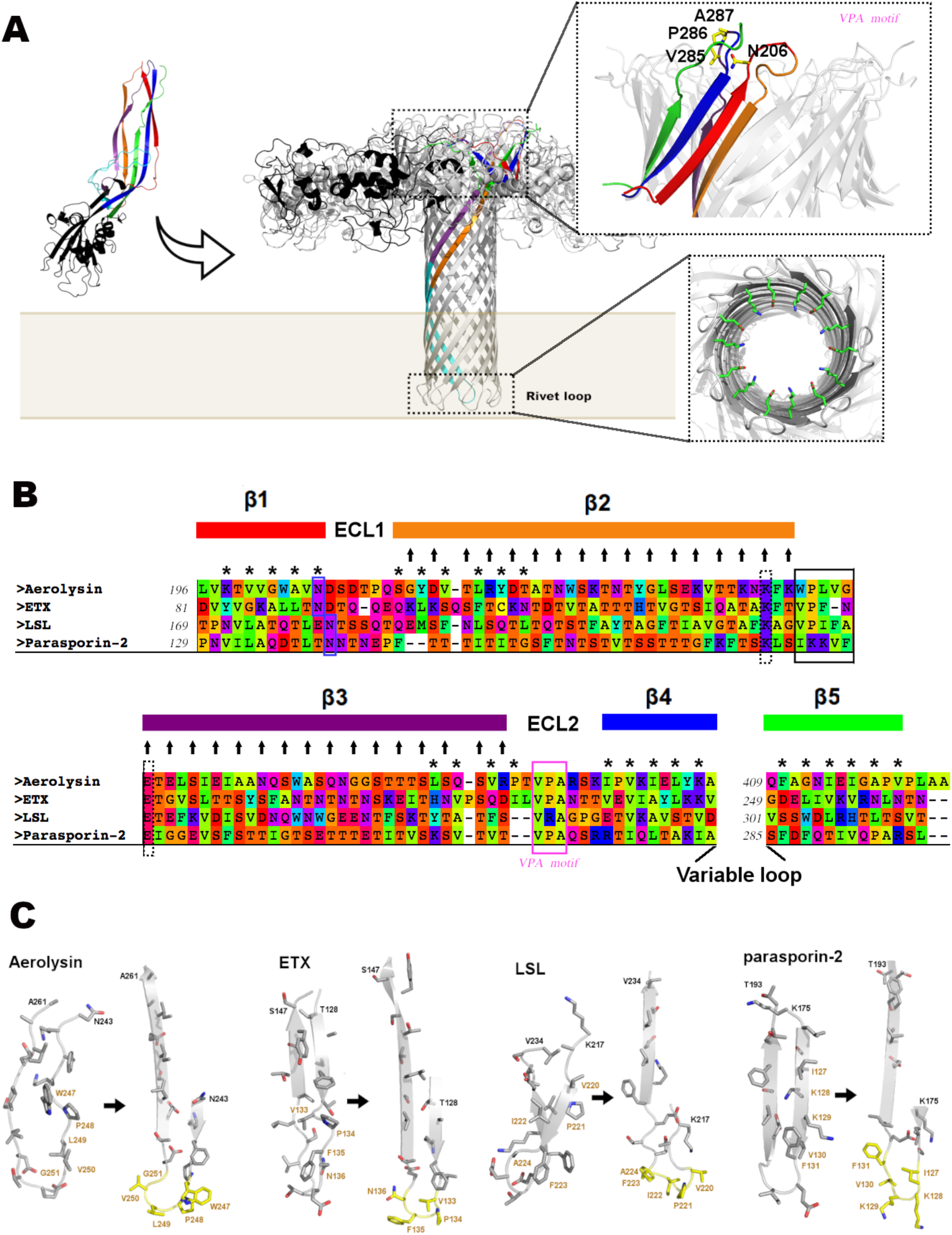
Analysis of the aerolysin-like models. **(A)** Structural architecture of the aerolysin-like family, shown on the aerolysin monomer (left) and the aerolysin pore (right). The following color scheme is used: membrane binding domain in black; the five β-strands of the pore-forming region colored in red (β1), orange(β2), purple (β3), blue (β4) and green (β3); and the stem loop between strands β2-3 in cyan. On the oligomers, only one protomer is colored. The upper close-up shows the conserved VPA motif (corresponding to the pink box in (**B**)) and the N206 of ECL1 (corresponding to the blue box in (**B**)). The lower close-up shows the two conserved charged residues (K&E) at the pore exit, corresponding to the dashed box in (**B**). This close-up is shown looking from the bottom of the pore. **(B)** Sequence alignment of the aerolysin-like pore-forming region used for creating the pore models. The β-strands are represented by solid bars using the same color scheme as in (**A**). The two loops at the top of the pore are named “extracellular loop 1” (ECL1) and “extracellular loop 2” (ECL2). The rivet loop is stressed by a black box, the conserved VPA motif by a magenta box, the conserved asparagine at the top of the barrel by a blue box, and the two conserved amino acids at the pore exit by a dashed box. The amino acids stabilizing the double β-barrel fold (those located between the two concentric rings) are marked with a star, while the amino acids which extend to the pore lumen are stressed by arrows. **(C)** For each toxin, the stem loop of the monomer structure (left), with the sliding of residue contacts that happen upon formation of the transmembrane pore, according to our models (right) is shown. The rivet loop is colored yellow.

Aerolysin and this family of proteins in general are receiving much attention because of their potential use in biotechnological applications. In fact, aerolysin is being studied for molecular sensing, comprising DNA, peptides, polysaccharide and polymers^4–7^. Biological nanopores when compared to inorganic solid-state nanopores possess a well-defined stoichiometry, and therefore a precise and stable pore diameter. Moreover, aerolysin pore presents other advantages, such as a long channel (∼10 nm) with a narrow diameter (∼1.0 nm), with two constrictions formed by arginine and lysine residues, which provide a slower translocation time of analytes compared to other commonly studied biological pores such as α-hemolysin^6,8^. Therefore, the existence of a potentially broad repertoire of aerolysin-like proteins holding similar properties opens a venue for improving and diversifying nanopore sensing capabilities. However, in spite of the conserved structural fold, the sequence identity in this family is very low. This complicates their sequence alignment, hindering a deep understanding of their pore structures and properties, as well as the possibility to rationalize their potential for further biotechnological applications.

Aerolysin presents a heptameric stoichiometry, while other proteins of this family have been found forming octamers (Dln1)^9^ or nonamers (Lysenin)^10,11^. In our previous study, we did not retrieve either Lysenin or Dln1 when searching for aerolysin-like sequences using the HHblits server^2^. On the other hand, three other aerolysin-like toxins with known monomer structure were recovered: epsilon toxin (ETX) from the anaerobic Gram-positive *Clostridium perfringens, Laetiporus sulphureus* lectin (LSL) and parasporin-2 from *Bacillus thuringiensis*. To note, ETX is known to form heptamers^12,13^, and oligomer assemblies of parasporin-2 in the membrane presented an estimated molecular weight of 200 KDa, indicating a potential hexameric/heptameric conformation^14^. Therefore, it can be speculated that sequence alignment algorithms manage to differentiate features defining stoichiometry, even if they are insufficient for properly aligning the amino acids in the pore region.

In an attempt to further understand the properties of aerolysin-like pores, we created models for the pore structure of three family members, the above-mentioned ETX, LSL and parasporin-2. To note, during the execution of this work, the structure of the ETX heptamer was released (PDB ID 6RB9) and was thus used to perform the following analysis. We selected those proteins for our study because they are the only ones recovered in our previous search which have the structure of the monomer published, and this has helped in creating the sequence alignment, as it will be explained later. Our models allowed us to further deepen into some conserved sequence/structural features of this family. Moreover, the protocol followed here to perform the sequence alignment could be applied to other family members, helping to unveil other potential candidates to improve and diversify the nanopore applications of this family.

To explore the potential translocation properties of these proteins, we performed molecular dynamics (MD) simulations in standard nanopore experimental setting (1.0 M KCl) and steered MD of ssDNA pass through the pore. Our results showed very different behavior for each protein pore, reconciling their different amino acid composition at the pore lumen. Furthermore, we performed ssDNA translocation experiments on ETX, the most promising of these pores. The results are consistent with our predictions, highlighting the possibilities of using the knowledge of the sequence and structure of these proteins to screen and explore new potential nanopore candidates. Interestingly, our experiments showed that ETX can form three different types of pores of different stoichiometry, where one conformation is able to translocate ssDNA. This unexpected behavior points towards the astonishing diversity produced by a structural pore motif shared by the aerolysin-like family of proteins.

## Results

### Refined sequence alignment to explore proteins as new potential nanopore candidates

Due to the low sequence identity among these proteins, automatic alignment tools fail to correctly align them, and therefore a manual alignment of aerolysin sequences was performed. While the pore-forming region of these toxins shares a similar fold, the same is not true for the membrane-binding domains. Therefore, these regions were not considered in our modeling approach. Looking at the structures of aerolysin in their different states (i.e., monomer, pre-pore, pore), the amino acid interactions that exist in the double β-barrel fold of the pore are already present in the monomer, even if the strands β2,3 and β1,4,5 are still not perfectly parallel and aligned as in the final pore structure. Consequently, we ensured to conserve the interactions observed in that region on the respective monomer structures of each toxin when building their pore models. Then, as no gaps could be introduced on the β-barrel section, the alignment of that region was straightforward, while some differences in sequence length were observed at the extracellular loops at the top of the pore. The curated sequence alignment of the pore-forming region of these toxins is shown in **Figure 1B**. Afterward, their pore models were built by homology modeling using the aerolysin structure as a template (PDB ID 5JZT)^15^, (see Methods section). Upon the release of the ETX structure, the analysis for this protein was done using PDB ID 6RB9^13^.

The polarity distribution of the pore structures showed the expected pattern, with the external surface of the pore split into two regions, the upper region that is mostly polar as it is exposed to the solvent, and a lower hydrophobic region, where the barrel is embedded in the lipid bilayer (**Supplementary Figure S2**). As expected, the pore lumen presents a hydrophilic character, being rich in small polar amino acids such as serine and threonine (arrows in **Figure 1B**). Of interest is the presence of a conserved pair of residues, one per inner strand, at the lower pore exit. The first is a positively charged lysine residue in strand β2 and the second is a negative glutamate residue at strand β3 (dashed square in **Figure 1B**, and lower close-up in **Figure 1A**). Those amino acids are not conserved for the lysenin pore (PDB ID 5GAQ)^11^.

In aerolysin, upon pore formation, the stem loop undergoes a sliding of residue contacts (7 residues) which results in the creation of a loop that has been suggested to act as a rivet helping to anchor the pore on the membrane^3,15^. Also, a sliding of 6, 8 and 7 residues are seen in the ETX structure, the parasporin-2 and LSL models, respectively (**Figure 1C**). Moreover, the polarity of the rivet loop is not conserved within the family, being highly hydrophobic in aerolysin, LSL and, to less extent, ETX, while positively charged in parasporin-2 (black square in **Figure 1B**); negative residues are observed for lysenin^11^. The rivet loop also exhibits differences in lengths among these proteins.

Our previous studies showed the conservation of motif at the top of the aerolysin pore (V285, P286, A287, name as VPA), as well as the double β-barrel region (upper close-up in **Figure 1A**). We observe this motif in aerolysin, ETX and LSL, and with some variations in LSL (magenta square in **Figure 1B**). This motif is also conserved in the more distant proteins lysenin and Dln-1, with small variations (IPP in lysenin and VPP in Dln-1, **Supplementary Figure S3**). Close by, an asparagine at the first top loop of the double β-barrel is also observed and conserved in all proteins within this family (upper close-up in **Figure 1A** and blue square in **Figure 1B**) ^2^. Again, lysenin and Dln-1 also present this amino acid (**Supplementary Figure S3**). The conservation of the residues of those loops agrees with the importance of the double β-barrel fold for folding and stability in this family. Then, we looked at the amino acids located between the two concentric β-strands, responsible therefore for stabilizing the fold (stars in **Figure 1B** and in **Supplementary Figure S3**). As described before, lysenin presents more polar, and even charged residues between the two rings, which has already been suggested as being related to the lower stability of lysenin pores compared to aerolysin^15^. On the other hand, the three aerolysin-like proteins studied here share with lysenin the hydrophobic pattern for the interactions stabilizing the double β-barrel fold.

### *In silico* analysis of pore sensing capabilities

The main characteristics controlling pore conductance are its length, diameter of the lumen and electrostatic properties. All the studied aerolysin-like proteins present similar lengths for the pore, while their diameter and electrostatics are however very different. In **Figure 2A-D**, a comparison of the radius of the pore for aerolysin and the three studied family members is shown. We can observe that aerolysin presents the narrowest constriction point of all proteins (∼0.58 nm of radius at R220, with a second constriction of ∼0.75 nm at K238). As we showed previously, these features are related with observed long dwell times of translocation and high sensitivity^8,16^. ETX and parasporin-2 possess only one constriction, at the pore entry in the case of ETX (∼0.75 nm, around K108) and at the pore exit for parasporin-2 (∼0.80 nm, around K158), whose lumen is constituted mainly by short side chains of threonine and serine amino acids. On the other hand, LSL does not show any narrower constriction feature, which could suggest some lower sensing capabilities, while on the other hand it may present a reduced signal to noise ratio.

**Figure 2.**
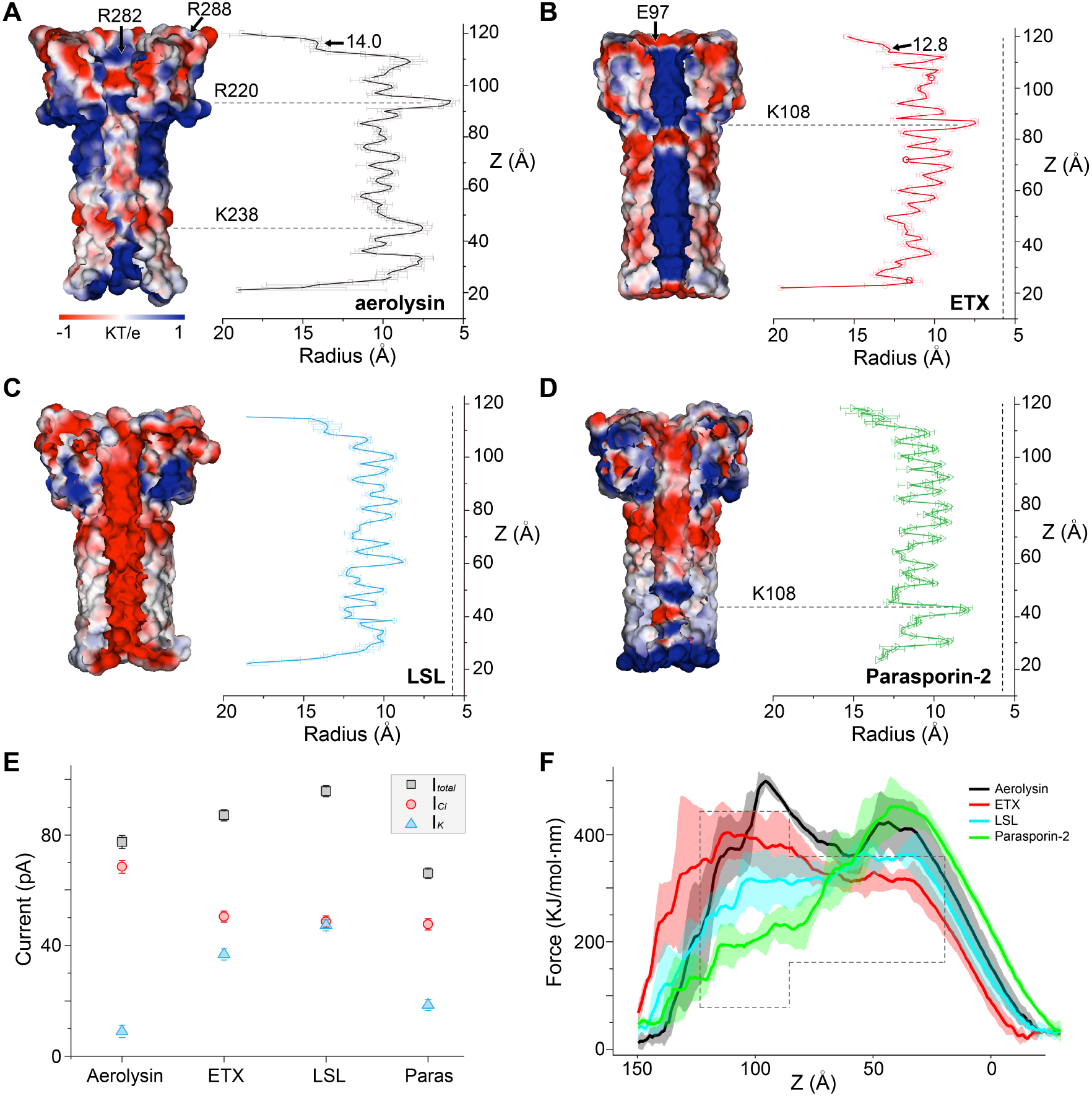
*In silico* analysis of aerolysin-like pore lumen electrostatics, radius, open-pore conductivity and ssDNA translocation. **(A-D)** Pore electrostatics and radius of the four proteins: aerolysin (**A**), ETX (**B**), LSL (**C**), and parasporin-2 (**D**). The pore electrostatics, on the left, were calculated by the PBEQ server, and the radius along the pore, on the right, was calculated from the MD simulations. The amino acids creating the main constriction points, when these exist, are shown for each protein, with a horizontal dashed line. In **B-D**, a vertical dashed line represents the narrowest constriction of aerolysin, for comparison. For aerolysin and ETX, the charged amino acids at the pore entry are shown, so the radius (in A) at the pore entry. **(E)** Open-pore current calculation by MD simulations of each pore at 150 mV, in 1.0 M solution of KCl electrolyte. The calculation of I_Cl_ (black) and I_K_ (red) is shown separately. (**F**) The force needed for dA_4_ to cross the pore at each section, as predicted by steered MD simulations. The position values correspond to the same as in panels **A-D**, and a schematic representation of a pore depicted in a dashed line has been depicted to help the illustration. The DNA enters from the cap, at the left of the plot, and moves towards the rivet (towards the right hand-side).

Concerning pore electrostatics, parasporin-2 shares with aerolysin its mainly neutral character, with moderate positive electrostatics at the pore exit (**Figure 2A** and **2B**). However, opposite to aerolysin, which possesses two rings of positive arginine at the pore entry, the upper region of parasporin-2 pore bears negative residues. On the other hand, ETX shows a strong positive potential all along the pore, while LSL has a strong negative potential (**Figure 2C** and **2D**)^17^. These differences in electrostatics could be exploited to extend the sensing demand to a wider range of molecules. Also, they will strongly influence the conductance of positive and negative ions. As so, in **Figure 2E**, we observe that, while parasporin-2 shares with aerolysin a conductivity controlled mainly by anions, in ETX and mainly in LSL both positive and negative ions are contributing similarly to the final conductivity. In total, higher conductivity values were estimated for ETX and LSL when compared to aerolysin, while parasporin-2 presented the lowest conductivity. In **Supplementary Figure S4**, the ion density of K^+^ and Cl^-^ along the pore is shown. As we discussed in a previous work^16^, when a negative voltage is applied in aerolysin, the density of negative ions is larger than that of positive ones, and they are mostly concentrated around the constriction of the pore. Following the negative electrostatics of LSL, a higher density of positive ions is observed, mostly around clusters of threonine residues (such as T195, T249 or T205) and the negative side chains of D231 and D235. Contrasting, however, with the positive character of the ETX lumen, the density of chloride ions was not higher than that of potassium. So as in parasporin-2, ion density was distributed evenly along the pore. Larger open pore current provides a bigger window of measurement, and therefore bigger differences between the detected molecules. This is interesting in cases when we need to identify a lot of different subunits, as for example the 20 amino acids of proteins. On the other hand, lower open pore current usually relates to higher sensitivity, as a molecule crossing the pore will have a significant effect in current blockage. Those two factors must be optimized for sensing purposes and having a spectra of aerolysin-like protein nanopores with the mentioned differences is therefore desirable.

In order to qualitatively explore the capabilities of the pores for molecular translocation, we performed steered MD of dA_4_ through each of them (**Figure 2F**). We could observe that both aerolysin and ETX pores present the highest crossing hindrance at the pore entry, while LSL and parasporin have it at the exit. Nonetheless, LSL does not show a clearly defined crossing point, coinciding with its uniform radius along the pore and lack of relevant constrictions (**Figure 2D)**. Parasporin-2, similarly to aerolysin, possesses a strong crossing barrier, with a force needed to translocate the molecule in MD simulations higher than 400 kJ mol^-1^ nm^-1^. Finally, compared to all other proteins, ETX presents the highest crossing hindrance at the very entrance of the pore (**Figure 2F)**.

### DNA translocation experiments in ETX

To further explore the differences in nanopore sensing predicted by MD, we selected the ETX pore to perform nanopore experiments to characterize its current response and capability to translocate ssDNA (i.e., dA_4_), as compared to the results obtained for aerolysin. When we incorporated the ETX into the planar lipid bilayer membrane (**Figure 3A**), we observed three different types of signatures, each bearing a different open pore current (OPC, **Figure 3B,C** and **Supplementary Figure S5**). The raw traces of OPC for the three types of pores at 100 mV is shown in **Figure 3B**. We observed a pore characterized by a small OPC of 20 pA (hereafter named type I), and two other pores with OPC of 50 pA and 60 pA, named type II and type III, respectively. Each type of pore has been observed to be stable, suggesting that ETX might be able to spontaneously assemble into nanopores bearing different sizes, and thus stoichiometry (**Figure 3C**). In fact, smaller oligomeric pores (type I) were already reported in a previous study^18^.

**Figure 3.**
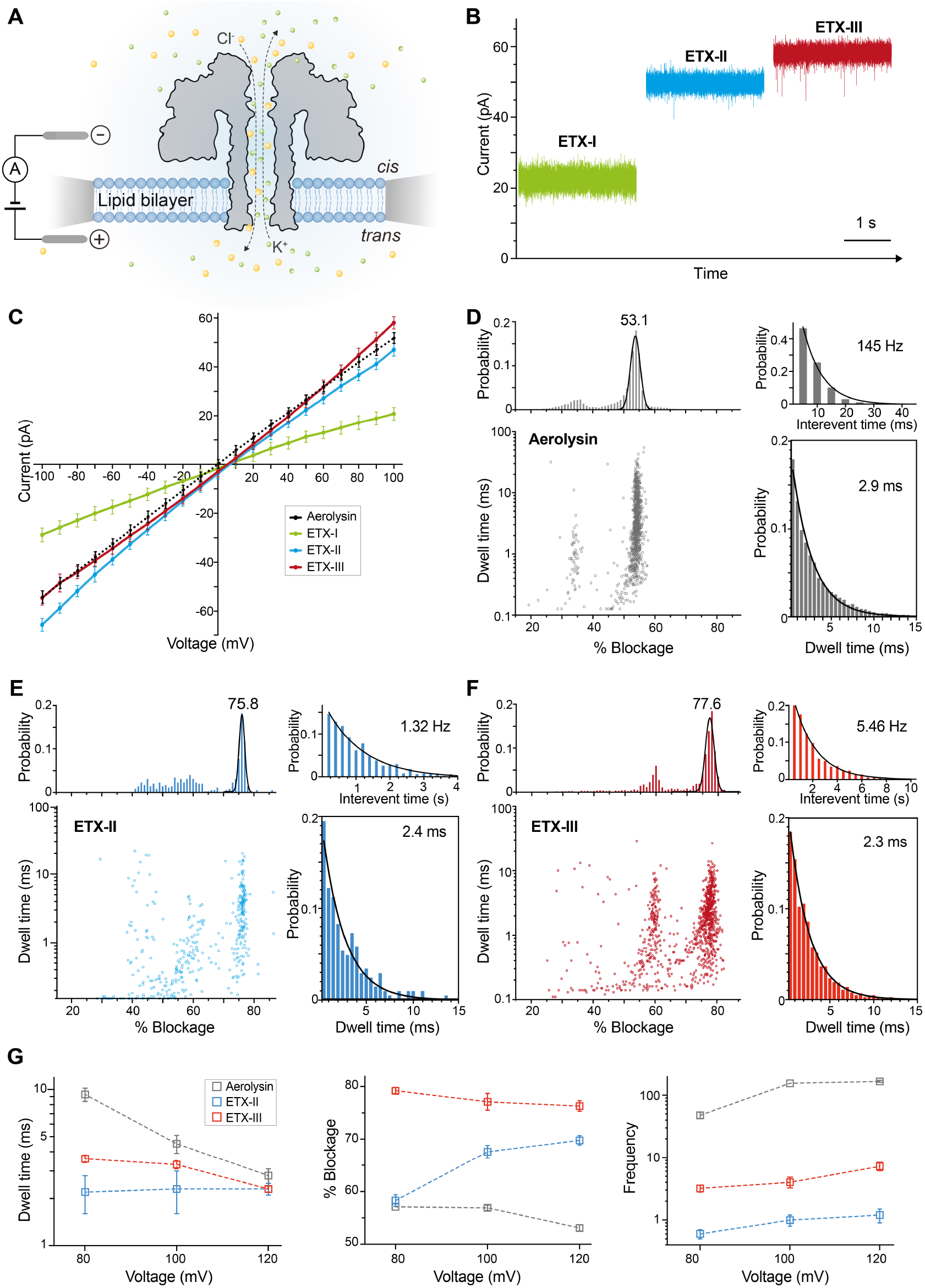
**(A)** A schematic representation of the ETX nanopore system, where the pore lumen is shown for the sake of clarity. **(B)** Typical raw traces of incorporation of the ETX pores at different oligomeric states into the lipid bilayer, where type I (green), type II (blue), and type III (red) are depicted. **(C)** IV curves of the three different ETX pores observed and WT aerolysin (dashed line). **(D-F)** The current histograms, the frequency distributions, the scatter plots, and the dwell time histograms and corresponding fitting of the events produced by the dA_4_ analyte through aerolysin **(D)**, ETX type II **(E)**, and ETX type III **(F)**. The OPC is represented as 0% by no pore blockage and at 100% by a pore fully blocked. For aerolysin WT and ETX type III, 1000 events were used for the scatter plot representation and 600 events were used for ETX type II. **(G)** The comparison of dwell time, % blockage, and event frequency of dA_4_ through WT aerolysin, ETX type II, and type III at different voltages. The values plotted here can be found in **Supplementary Table S1**.

Having studied the pore formation and OPC, we analyzed dA_4_ events through ETX pores and compared them with wild-type aerolysin (**Figure 3**). We investigate and analyze if the events produced by dA_4_ represented a transport process through the pore (translocation events) or if they were interacting with the pore without any transport through it (bumping and/or trapping events). The difference between the two types of events is that upon increasing applied voltages, a decrease in dwell time is associated with a transport process, since an increased potential difference gives rise to a stronger driving force, therefore dragging the analyte faster through the pore^6,19^. In contrast, if the analyte is not transported along the pore, it will interact with the pore, and the dwell time will increase upon increased applied voltage. As shown in **Figure 3G** left panel and **Supplementary Table S1**, the dwell time for aerolysin and ETX pore III is decreasing significantly upon increasing voltage, implying that ssDNA is transported across the pore. Whereas for the ETX type II, the dwell time does not change significantly over voltage increase, implying that the dA_4_ is not transported in ETX type II, likely due the too narrow diameter of this pore state. We also analyzed how the current blockage (i.e., 0% for no pore blockage and 100% for a pore fully blocked) is affected in the different toxins by DNA upon voltage increase (**Figure 3G** middle panel). By increasing voltage, the current blockage generated by the DNA molecule going through aerolysin is slightly decreasing (as previously observed in the detection of negatively charged peptides in aerolysin^20^) and ETX type III is following the same trend. Whereas for ETX type II, the current blockage is increasing and as shown in **Supplementary Figure S6**, most of the events did not trigger a large blockage, but only small current blockage (a region comprising 0-25% current blockage), further implying that the DNA molecule is not transported in ETX type II but interacting with the pore.

The event frequency observed for ssDNA captured by ETX type II and III are very low compared to aerolysin (**Figure 3G**, right panel). This could be related to the bigger force hindrance observed previously in steered MD simulations at the pore entry for ETX (**Figure 2F**). Looking at the residues located in this region for both aerolysin and ETX proteins (**Figures 2A/B**), we can observe that in ETX, a negatively charged amino acid at the top of the pore, E97, forms a ring of negative charges, therefore creating a repulsive force towards the DNA molecule and impeding the translocation of DNA. On the contrary, aerolysin presents two arginines (R288, R282) at the pore entry, which may help capturing DNA instead. Similarly, previous works on aerolysin have shown that the mutation of one of those arginines (i.e., R282A) abolished DNA captures^16^. Moreover, comparing the radius of ETX and aerolysin at the very pore entry (**Figures 2A/B** and **Supplementary Figure S7**), we observed that ETX is smaller. We think that the combination of the negative charge distribution and smaller radius of ETX at the pore entry compared to aerolysin strongly reduces the capture ability of negatively charged DNA (**Figure 3G**).

Altogether, we observed faster dwell times in ETX compared to aerolysin when the applied voltages were higher than 100 mV (**Figure 3G**), coinciding with its bigger diameter and smaller maximum force predicted by steered MD (**Figures 2F**). Surprisingly, the current blockage by DNA is bigger in ETX. This could be related to its stronger positive character along the pore. Together with the presence of E97 at the top of the pore, ETX could be studied for sensing more positive molecules. Moreover, the larger current blockade could be an interesting feature since it suggests that the sensitivity of molecules such as DNA, peptides, or other (bio)polymers could be enhanced compared to aerolysin.

## Conclusion

Biological nanopore sensing has attracted growing attention during the last few decades because of the possibility of sequencing long DNA in a cheaper and faster way than ever before. Therefore, this technology has been studied for a broader range of applications. The discovery of novel proteins that could be developed into new more sensitive nanopores that can further expand their fields of use is therefore highly desirable. However, the structure of β-pore-forming proteins is difficult to obtain, even to model, making it difficult to introduce novel nanopores in the field. In this work, we used the knowledge of the monomer structure of proteins of the aerolysin family to build 3D models of their oligomeric pores. This allowed us to predict some properties related to translocation, such as steric and electrostatics, and to perform predictions of their OPC and the hindrance for DNA translocation. The experimental measurements performed for one of them, ETX, confirmed our predictions, validating the use of *in silico* calculations for studying mechanisms related to nanopore sensing.

Interestingly, we discovered an unexpected and still unknown behavior for ETX, which formed three types of pores (**Figure 3A**), with increasing conductance with larger conductance pore being more prone to form (ETX type III with 44% occurrence, **Supplementary Figure S5**). Unfortunately, due to the lack of structural information, it is hard to conclude which type corresponds to which oligomeric state. Comparing the experimental data with our modeling predictions, we think that ETX type III, showing a OPC similar to aerolysin, would represent the expected heptameric state, but an alternative octameric state cannot be ruled out. This conformation is also the only one able to translocate short ssDNA, as aerolysin does. ETX presents a very low frequency of translocation, which seems to be related to a small pore entry and the presence of a negative ring of residues (i.e, E97) at the top, probably repelling negatively charged molecules such as DNA. However, this could be an advantage for sensing positive or even neutral molecules. Moreover, ETX showed a larger current blockage than aerolysin and only one constriction region, properties compatible for increasing the sensitivity towards the sensing of smaller molecules, DNA, and peptides. Altogether, these results open new possibilities for molecular sensing using the different conformations of the ETX pores, for improving experimental conditions for the selection of one type of pore or for the development of novel ETX mutants.

## Methods

### Modeling of pore structures

Only the β-barrel region was considered for the models, without the membrane-binding domains. This corresponds to aerolysin residues 24–195, 301–408, and 425–447. The sequence alignment was performed manually based on the structural superposition of the monomer structures: aerolysin (PDB ID 1PRE)^21^, ETX (PDB ID 1UYJ)^22^, LSL (PDB ID 1W3A)^23^ and parasporin-2 (PDB ID 2ZTB)^24^. Some conditions had to be respected for the alignment: i) ensure the preservation in the pore models of the amino-acid contacts observed on the monomers between strands β2,3 and strands β1,4,5, which will form the double β-barrel fold; ii) gaps are only allowed in the loops. Then, a model for a heptamer pore was built with the Modeller program v9.11^25^, using the aerolysin pore structure as a template (PDB ID 5JZH)^15^, and following the sequence alignments created above. Upon release of the ETX pore structure (PDB ID 6RB9)^13^, we were able to observe that our proposed alignment (and therefore model) agreed with the real structure for this protein (a structural comparison is provided in **Supplementary Figure S8**).

### Molecular Dynamics

Molecular dynamics of the pore models on a lipidic membrane was conducted with the Gromacs software version 2018^26^. To note, in the case of ETX, the published structure was used instead of our model. First, the membrane position was estimated by the PPM server^27^, and afterwards a ∼9 × 9 nm^2^ membrane bilayer was modeled by 1-palmitoyl-2-oleoylphosphatidylcholine (POPC) lipids using the CHARMM-gui server^28^. These systems were then solvated in a 1 M KCl water box with initial dimensions ∼9 × 9 × 15 nm^3^. All MD simulations were run using the CHARMM36 force field^29^, the SHAKE algorithm on all the bonds between hydrogen and heavy atoms, and Particle-Mesh Ewald, treating the electrostatic interactions in periodic boundary conditions. Each system was first minimized using the steepest descent algorithm, and afterwards equilibrated using a similar protocol that suggested by the CHARMM-gui server, with 6 steps restraints in proteins and lipids were gradually reduced to zero: two steps of 50 ps each on the NVT ensemble (1 fs of time step), and four steps on the NPT ensemble, the first one (50 ps) with a time step of 1 fs and the last three steps (100 ps each) with a time step of 2 fs. This time step was used for the rest of the simulations. A temperature of 22 °C was controlled with the Nose-Hoover thermostat and the Parrinello-Rahman method was used for semi-isotropic pressure coupling. After equilibration, we followed with an unrestrained simulation for 250 ns, applying an electric field corresponding to 150 mV. The RMSD plots during the equilibration and production phases for all studied systems can be found in **Supplementary Figure S9**. Steered MD was used to explore the DNA crossing hindrance along the pores, using an umbrella biasing potential, based on direction-periodic geometry and using as reaction coordinate along the z axis, with the pores aligned to it. A dA_4_ molecule was placed above the aerolysin pore, aligned to the z-axis and with its 5’ terminus pointing down, located around 20 Å distance to the pore entry. Then, the structures of ETX, LSL and parasporin pores were superposed to the starting conformation of aerolyin/DNA, to ensure that the distance between pores and DNA was the same for all proteins at the start of the simulation, and to help compare the observed forces. A harmonic biasing force (with a spring constant of 100 kJ mol^−1^ nm^−2^) was applied to the spring connecting the center of mass of the 5′ nucleotide, and the center of mass of the α-carbons of the seven conserved Lys residues, located at the pore exit, at a constant velocity of 0.004 nm ps^−1^. Three replicas were performed for each protein with different random seeds for initial velocities, using the same initial structures.

### *In silico* analysis of pore properties

Electrostatic calculations were performed with the PBEQ server^30^, which considers the presence of a lipid bilayer. As input for the calculations, it was used the protein conformation closest to the average of each production run (for aerolysin, the structure with PDB ID 5JZH was used), oriented with the PPM server. The calculation was performed with the server default parameters.

To calculate the OPC, so as for the ion density map, all MD trajectories were cut to the same length from 50 to 100 ns so the analysis was performed on the equilibrated part of the trajectory. A VMD script was used to calculate instantaneous ionic currents as a sum of shifts of ions every 10 ps as described in previous studies^16^. Then the cumulative current was calculated, and a linear regression was used to extract the average value of the current. For the ion density calculation, all trajectories were aligned to the same reference structure to minimize RMSD of the beta-barrel part of the pore, using the VMD RMSD Trajectory Tool. This procedure allows to minimize errors related to rotation of the pore and to consistently compare data from different simulations. The calculation of density for water, Cl^-^ and K^+^ was performed with the VMD plugin volmap density, at a resolution of 1 Å every 100 ps, and results were averaged to obtain mean and deviation. Then we used the method described in the past study^31^ to calculate the radius of the pore cavity occupied by water and the average density of Cl^-^ and K^+^ inside the cavity. The location of the pore axis was determined by the center of mass of the protein, then the volume was sliced into layers of 1 Å and the radius was found by an iterative process: first, the radius was assigned to be 1 Å, and later, it was increased by 0.5 Å while the ratio of water/non-water atoms inside the added ring was larger than 25%.

### ETX expression and purification

The ETX (Uniprot sequence Q57398, residues 47-328) expression plasmid was a generous gift from Prof. Monika Bokori-Brown and carries the gene in the pHIS-Parallel1 vector, with an N-terminal Hexa-histidine tag, followed by a TEV (Tobacco etch virus) cleavage site. The plasmid was transformed into Rosetta (DE3) cells (Promega). Protein expression was induced by the addition of 0.25 mM isopropyl ®-D-1-thiogalactopyranoside (IPTG) when the cells reached an optical density of 0.6 and subsequent growth overnight at 16°C. Cell pellets were resuspended in lysis buffer (0.5 M NaCl, 20 mM Na_3_PO_4_ pH 7.4 and cOmplete™ Protease Inhibitor Cocktail (Roche) and then lysed using sonication. The resulting suspension was then centrifuged (13,000 rpm for 35 min at 4°C) and the supernatant was applied to a HisTrap HP column (GE Healthcare) previously equilibrated with the lysis buffer. The protein was eluted with a continuous gradient over 40 column volumes of elution buffer (0.5 M NaCl, 20 mM Na_3_PO_4_ pH 7.4, 500mM Imidazole.). Subsequently, pure fractions were buffer-exchanged into the final buffer (20 mM Tris-HCl, pH 8, and 150 mM NaCl) using a HiPrep Desalting column (GE Healthcare) and stored at -20°C in the final buffer. Protein table integrity was checked by mass spectrometry (results not shown).

### Single-channel recording experiments

Phospholipid of 1,2-diphytanoyl-sn-glycero-3-phosphocholine powder (Avanti Polar Lipids Inc., Alabaster, AL, USA) was dissolved in octane (Sigma-Aldrich Chemie GmbH, Buchs, Switzerland) to a final concentration of 8 mg/mL. Purified aerolysin and ETX were activated as previously described^13,16^. Briefly, the toxin was diluted to the concentration of 0.2 μg/ml and then incubated at 4°C with Trypsin-agarose (Sigma-Aldrich Chemie GmbH, Buchs, SG Switzerland) for 2 hours to activate the toxin for oligomerization. The solution was centrifuged (10,000*g*, 4°C, 10min) to remove the trypsin-agarose beads. Activated toxins were aliquoted and kept at -20°C.

Nanopore single-channel recording experiments were performed on an Orbit Mini instrument equipped with a temperature control addon (Nanion, Munich, Germany). Phospholipid membranes were formed across a MECA 4 recording chip that contains 4 circular microcavities (size 50µm diameter) in a highly inert polymer. Each cavity contains an individual integrated Ag/AgCl microelectrode and can record four artificial lipid bilayers in parallel. The buffer (10mM Tris, 1mM EDTA, 1M KCl, pH 7.4), the concentration of analyte (6 µM) and the temperature were kept constant, and the temperature was set to 25°C for all experiments. Activated toxins were added and insertion of pore and I-V curves were recorded for each toxin. Once a stable baseline has been recorded, the dA_4_ DNA (Integrated DNA technology, USA) analyte was added into the *cis* chamber, and the events were recorded in Elements Data Reader (Elements srl, Italy) and further analyzed by using Clampfit (Axon, Molecular device) and Nanolyzer software (Northern Nanopore Instruments). Results, fitting, and graphs were produced in Prism (GraphPad, v9) and figures and tables were generated in Adobe Illustrator 2022 (Adobe).

## Supporting information

This PDF file includes: Supplementary Figures 1 to 9 Supplementary Table S1

## Notes

### Competing Interest Statement

The authors have declared no competing interest.

