## Supplementary material for "Exploring new nanopore sensors from the aerolysin family": This PDF file includes: Supplementary Figures 1 to 9 Supplementary Table S1

Supplementary Figures 1 to 9
Supplementary Table S1

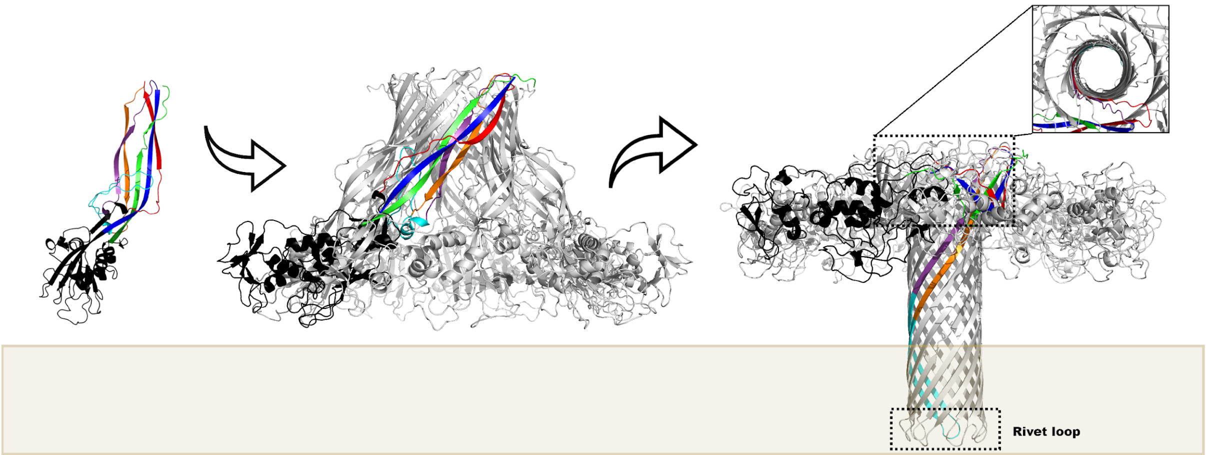


**<u>Supplementary Figure 1.</u> Aerolysin structures and its pore-forming mechanism**. From left to right, aerolysin monomer (PDB ID 1PRE)[11], oligomerization into the pre-pore state (PDB ID 5JZH),[12] and formation of the heptameric pore (PDB ID 5JZT)[12] in the membrane (represented as a brown shadow). The following color scheme is used: membrane binding domain in black; the five β-strands of the pore-forming region colored in red (β1), orange (β2), purple (β3), blue (β4) and green (β3); and the stem loop between strands β2-3 in cyan. On the oligomers, only one protomer is colored. In the closeup, the double β-barrel fold is shown. The rivet loop formed after residue sliding of the stem loop is stressed.

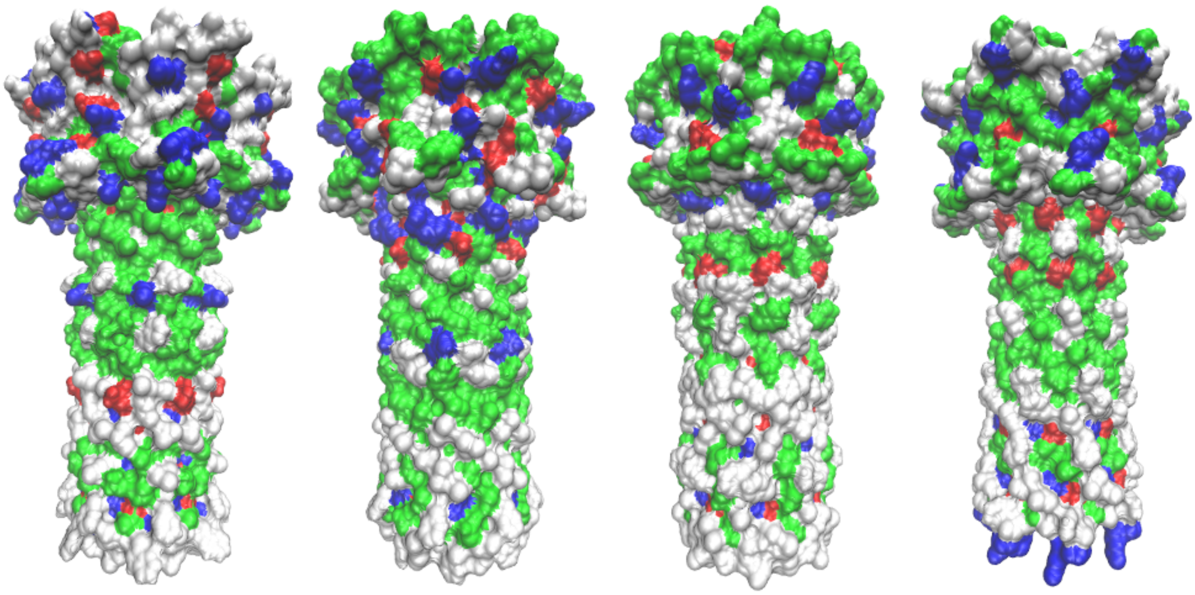

**<u>Supplementary Figure S2</u>. Molecular surface of the transmembrane and DBB regions of the aerolysin-like proteins**. The surfaces are colored by polar characters. From left to right: aerolysin, ETX, LSL and parasporin-2. Blue means positively charged residues while red denotes negatively charged ones. Green color is for polar neutral amino acids, and white is for hydrophobic ones.

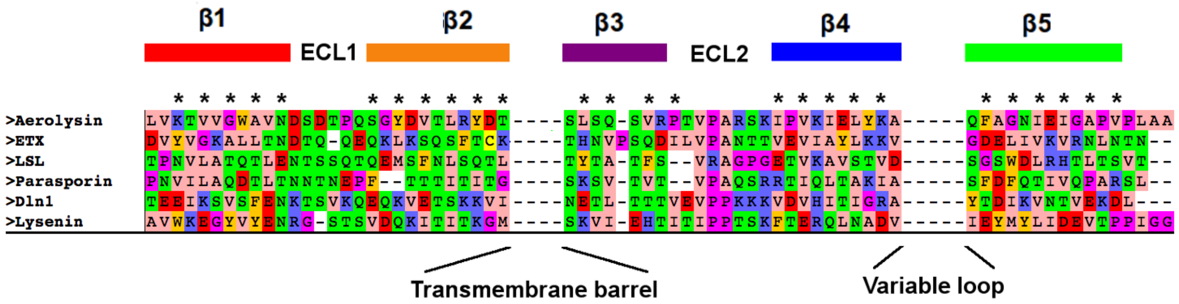


**Supplementary Figure S3. Sequence alignment of the double-beta barrel region of aerolysin-like proteins with Dln-1 and Lysenin.** The β-strands are marked by bars using the same color scheme as in Figure 1. The two loops at the top of the pore are named “extracellular loop 1” (ECL1) and “extracellular loop 2” (ECL2). The conserved VPA motif and its neighbor asparagine are shown by a red mark. The amino acids stabilizing the double β-barrel fold (those located between the two concentric rings) are marked with a star. The position of transmembrane and membrane-domain is marked on the sequence.

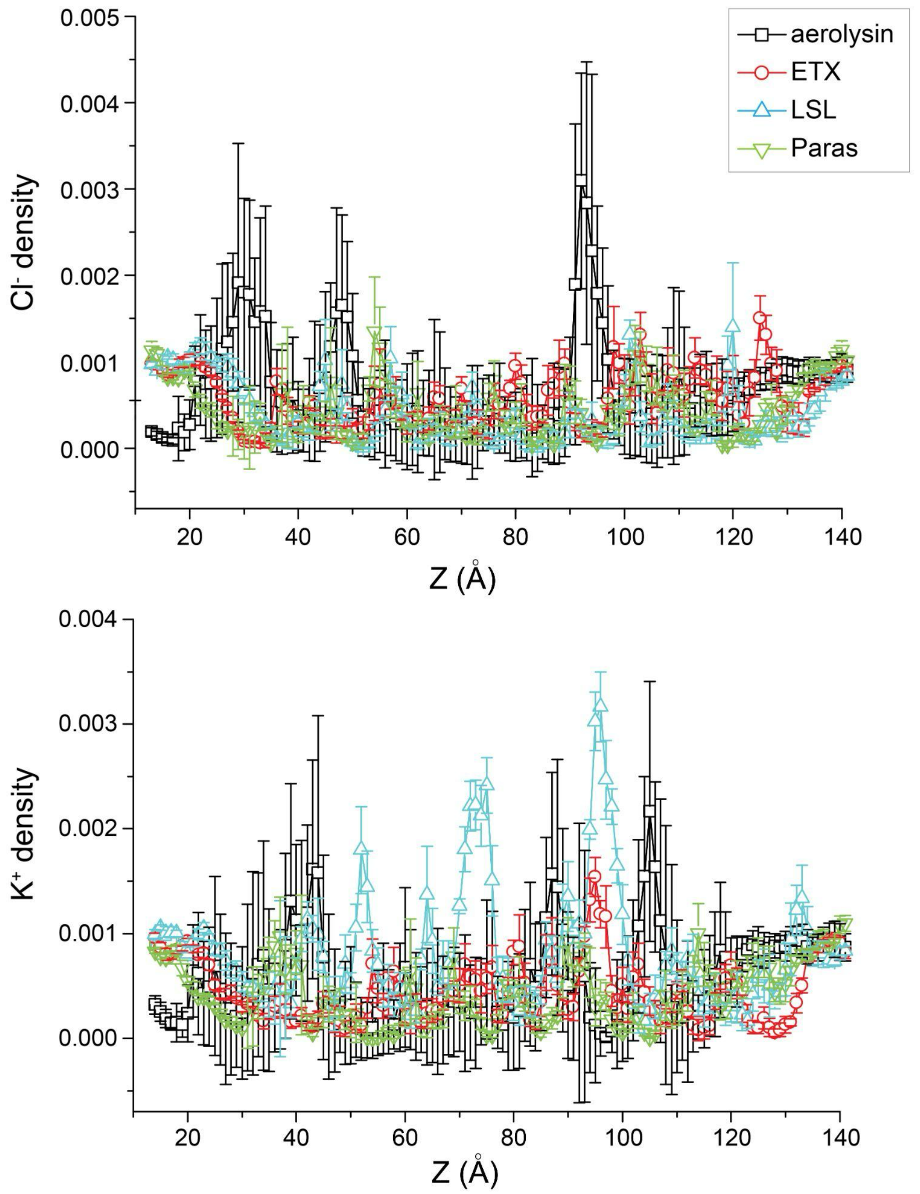


**<u>Supplementary Figure S4.</u> The ion density maps of aerolysin and aerolysin-like proteins.** The z axis is shown from the pore exit (rivet) to the pore entry (cap). The averaged density of $Cl^-$ (up) and $K^+$ (bottom) at +150 mV along the z axis of the pore, as defined by the local radii profile during 250 ns MD simulation.

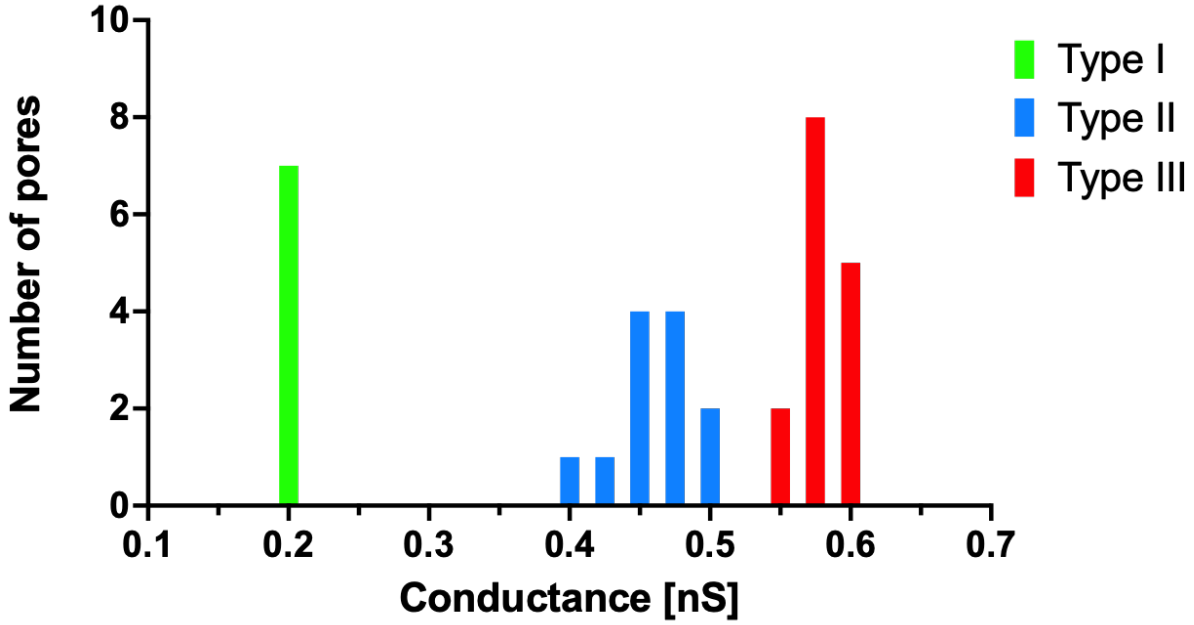


**Supplementary Figure S5.** Conductance of ETX from the different types of pores observed experimentally. Occurrence is as follow: Type III is prominent (44.1%), then Type II (35.3%) and finally Type I (20.6%)

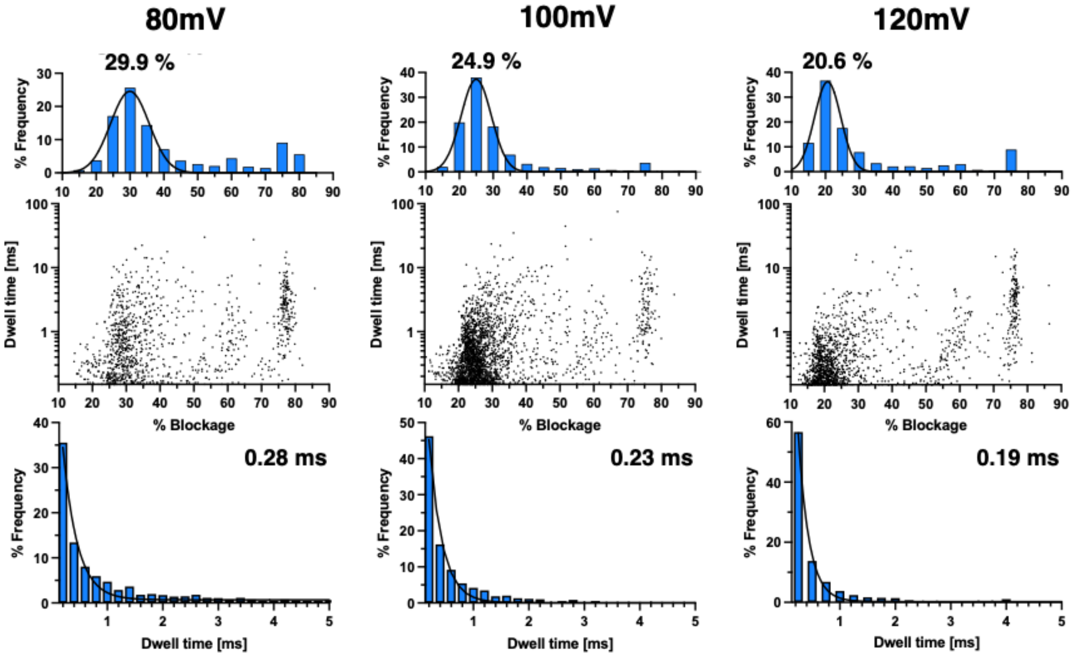


**Supplementary Figure S6.** Scatter plots showing the events below 30% at different voltages for ETX pore type II.

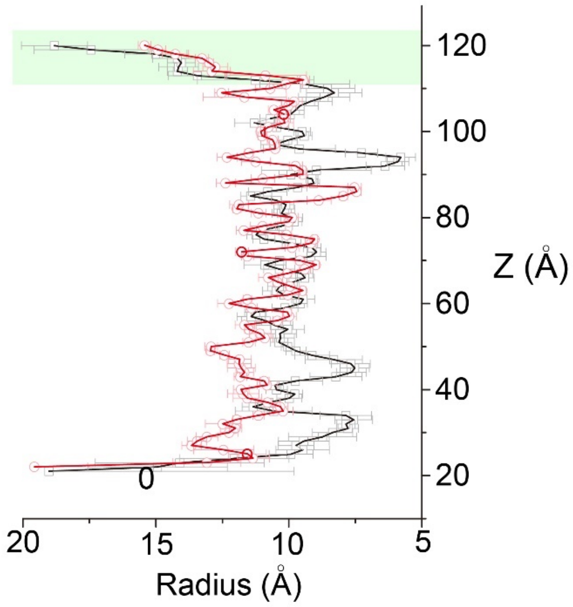


**<u>Supplementary Figure S7</u>.** Superposition of the measurements of the radius of aerolysin (black) and ETX (red) pore lumen, corresponding to **Figures 2A-B**.

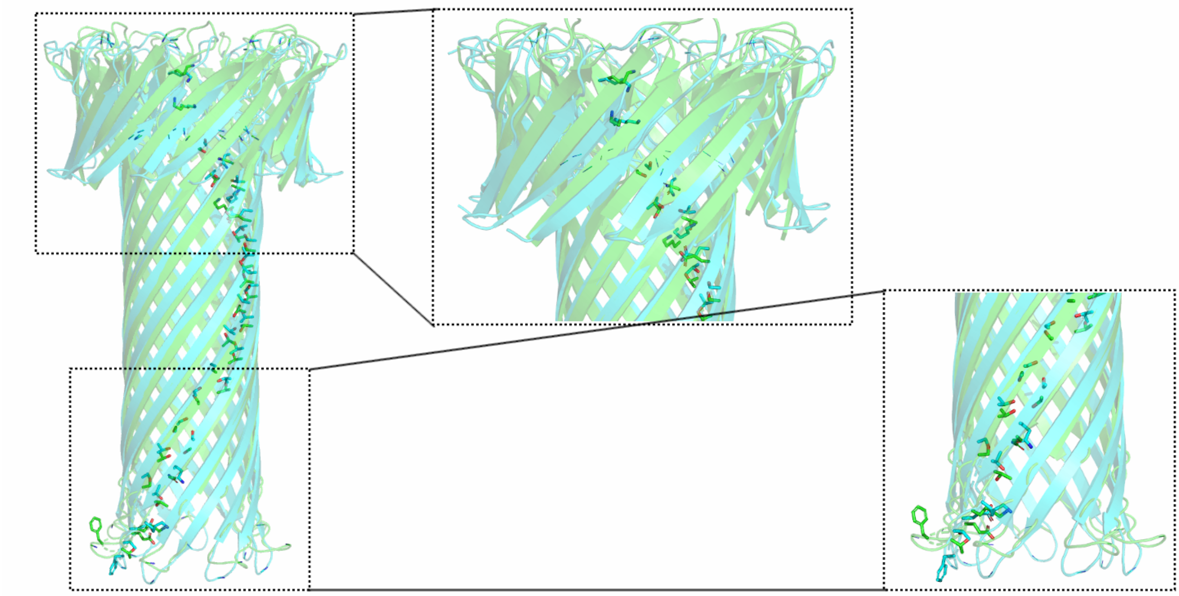

**<u>Supplementary Figure S8</u>.** Superposition of our ETX model (cyan) and the experimental structure with PDB ID 6RB9 (green). The residues in the lumen are shown, but only for one chain to help visualization. For visualization, two close-ups of the cap and the membrane region are shown. As explained in the methods section, the membrane-binding domains were not included in this study.

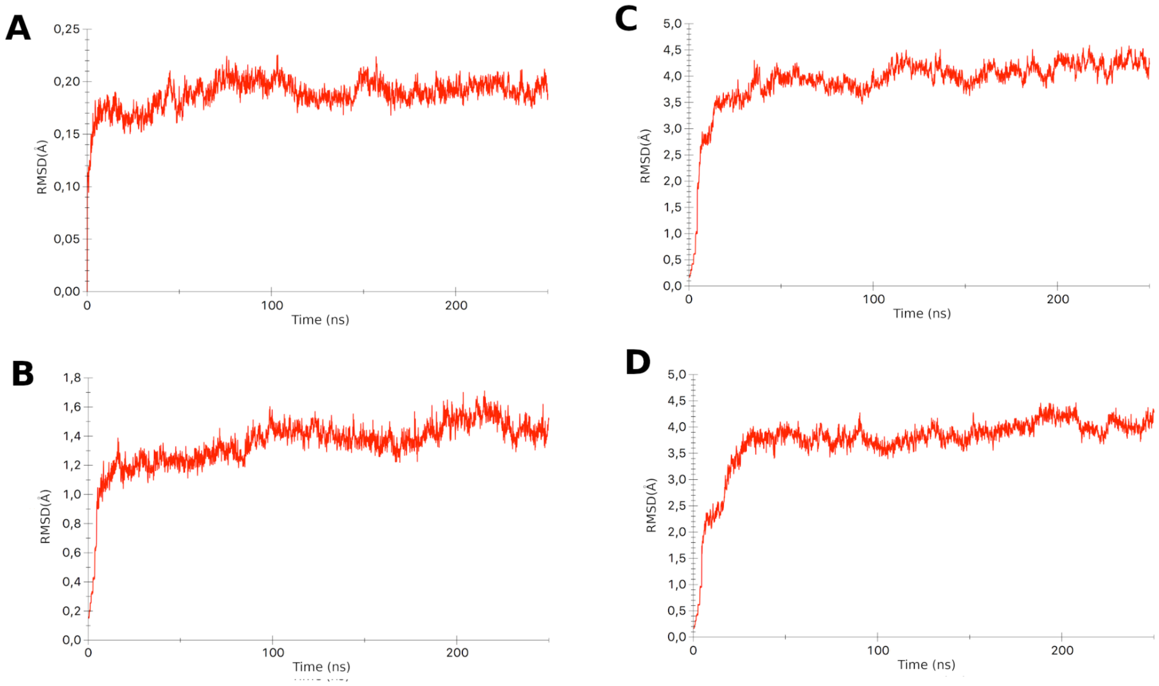


<u>**Supplementary Figure S9**</u>**. Root Mean Square Deviation (RMSD) of the backbone residues of the pore structures along MD simulations.** (**A**) aerolysin, (**B**) ETX, (**C**) LSL, (**D**) parasporin-2.

| Voltage [mV] | Dwell time [ms] | | | % Blockage | | | Frequency (Hz) | | |
|---|---|---|---|---|---|---|---|---|---|
| | AeL WT | ETX Type II | ETX Type III | AeL WT | ETX Type II | ETX Type III | AeL WT | ETX Type II | ETX Type III |
| 80 | 9.3±0.9 | 2.2±0.6 | 3.6±0.2 | 57.1±0.3 | 58.3±1.1 | 79.2±0.5 | 51.8±3.8 | 0.6±0.1 | 3.2±0.4 |
| 100 | 4.5±0.6 | 2.3±0.7 | 3.3±0.2 | 56.9±0.5 | 67.6±1.2 | 77.1±1.6 | 104.4±7.3 | 1.0±0.2 | 4.0±0.5 |
| 120 | 2.8±0.3 | 2.4±0.2 | 2.3±0.1 | 53.1±0.6 | 69.8±0.9 | 76.3±1 | 165.9±9.1 | 1.2±0.3 | 7.2±0.7 |

<u>**Supplementary Table S1.**</u> **Values plotted in the chart provided in Figure 3.** The error is calculated from 3 different pores at the given voltage. Data below 50% blockage current has not been considered for the calculation of the dwell time and the event frequency.
